# Myocardin ablation in a cardiac-renal rat model

**DOI:** 10.1101/470708

**Authors:** Anupam Mittal, Santanu Rana, Rajni Sharma, Akhilesh Kumar, Rishikesh Prasad, Satish K Raut, Sagartirtha Sarkar, Uma Nahar Saikia, Ajay Bahl, Perundurai S Dhandapany, Madhu Khullar

## Abstract

Cardiorenal syndrome is defined by primary heart failure conditions influencing or leading to renal injury or dysfunction. Dilated cardiomyopathy (DCM) is a major co-existing form of heart failure (HF) with renal diseases. Myocardin (MYOCD), a cardiac-specific co-activator of serum response factor (SRF), is increased in DCM mice and patient cardiac tissues and plays a crucial role in the pathophysiology of DCM. Inhibiting the increased MYOCD has shown to be partially rescuing the DCM mice phenotype. However, expression levels of MYOCD in the cardiac tissues of the cardiorenal syndromic patients and the beneficial effect of inhibiting MYOCD in a cardiorenal syndrome model remains to be explored.

Here, we analyzed the expression levels of MYOCD in the DCM patients with and without renal diseases. We also explored, whether cardiac specific silencing of MYOCD expression could ameliorate the cardiac remodeling and improve cardiac function in a renal artery ligated rat model (RAL). We observed an increase in MYOCD levels in the endomyocardial biopsies of DCM patients associated with renal failure compared to DCM alone. Silencing of MYOCD in RAL rats by a cardiac homing peptide conjugated MYOCD siRNA resulted in attenuation of cardiac hypertrophy, fibrosis and restoration of the left ventricular functions.

Our data suggest hyper-activation of MYOCD in the pathogenesis of the cardiorenal failure cases. Also, MYOCD silencing showed beneficial effects by rescuing cardiac hypertrophy, fibrosis, size and function in a cardiorenal rat model.

## Introduction

DCM is a major cause of HF,^1^ accounting for nearly 1/3rd of total cases. Many of these patients subsequently display kidney dysfunction or injury leading to cardiorenal syndrome. More than half of the heart failure patients show renal diseases. Co-existence of cardiac and renal dysfunction in the patients increases the mortality significantly compared to cardiac or renal disease alone patients.

Various molecular pathways including Renin-angiotensin-aldosterone system (RAAS) are shown to be influencing the cardiorenal syndrome. Notably, circulating Ang II (an essential component of RAAS) affects cardiac function by, increasing systemic arteriolar vasoconstriction, vascular resistance, and cardiac afterload through AT1 receptor-mediated endothelial dysfunction ^2^.

Ang II has been shown to induce MYOCD under hypoxic condition^3^. MYOCD is a cardiac-specific transcriptional co-activator present in cardiomyocytes and smooth muscle cells. MYOCD is involved in heart development and cardiomyocyte differentiation^4,5^. Also, MYOCD is required for maintenance of structural integrity, cardiomyocyte survival, and heart function^5–7^. Besides, cardiomyocytes and smooth muscle cells MYOCD expression has been detected in fibroblasts^8^. MYOCD has been shown to promote fibroblast to myofibroblast differentiation and to inhibit cell proliferation ^9^. Forced expression of MYOCD in fibroblasts induces cardio-myogenic properties alone^8^ and/or in combination with other factors^10^. Transforming growth factor (TGF-β) was shown to induce MYOCD expression in fibroblasts and vice-versa^9^. TGF-β induction of MYOCD expression in the infarcted heart may have a potential function in fibroblast-to-myofibroblast transition, similar to Myocardin related transcription factor (MRTF)-A and MRTF-B which have been shown to be key regulator in fibroblast to myofibroblast differentiation induced by TGF-β1^11^. Further, deletion of MYOCD gene in the adult murine heart resulted in dilated cardiomyopathy, and rapid death due to heart failure^5^. Upregulation of MYOCD expression has been shown in cardiac hypertrophy^3,12,13^ and MYOCD overexpression in mouse cardiomyocytes resulted in activation of genes associated with cardiac hypertrophy^12^. Increased cardiac MYOCD expression has been reported in various cardiac ailments including DCM patients with end-stage HF ^14,15^. MYOCD has been shown to be a pro-hypertrophic factor in cardiac remodeling induced in multiple models^3,12,13^. However, there is no report so far, suggesting the role of MYOCD in cardiorenal syndrome.

In the present study, we analyzed the cardiac-specific expression of MYOCD in DCM patients with renal disease and DCM alone cases. The results showed the MYOCD is overexpressed in the DCM patients with renal disease compared to DCM alone cases. In addition, the effects of cardiac-specific silencing of MYOCD was explored in a cardiac renal syndrome rat model. The cardiac-specific silencing of MYOCD in rats decreased the expression of upregulated hypertrophic and fibrotic genes leading to restoration of left ventricular function.

## Material and Methods

### Study Population

Thirty consecutive biopsies were taken from left ventricle region from idiopathic DCM (IDCM) patients, attending Cardiology Clinic at the Department of Cardiology, Postgraduate Institute of Medical Education and Research, Chandigarh, India between Jan 2011-2014. Inclusion criteria for recruitment of DCM patients, diagnosed after echocardiography, defined by left ventricular ejection fraction (LVEF) ≤ 40% and chronic mild to severe HF (NYHA functional class II to IV). All patients underwent left cardiac catheterization and coronary angiography before their inclusion in the study. Exclusion criteria were: the presence of significant coronary artery disease defined as lumen stenosis in >50% of any coronary artery, severe primary valve disease, uncontrolled systemic, hypertension, hypertrophic or restrictive cardiomyopathy, chronic systemic disease like myocarditis, thyrotoxicosis, HIV disease and drug abuse. All recruited IDCM subjects were on optimal medication, angiotensin-converting enzyme inhibitors, and beta-blockers but had persistently low LVEF despite drug regime at the time of biopsy. Endomyocardial biopsy from left ventricle region (n=14) taken from subjects undergoing surgery for ventricular septal defect (VSD), served as controls. The VSD patients recruited in the study have normal LVEF with no right or left ventricular dysfunction. The study was approved by the Institutional Ethics Committee (8443-PG-1TRg-10/4497), Postgraduate Institute of Medical Education and Research, Chandigarh and written informed consent was taken from all patients for participation in the study. We confirm that all experiments were performed in accordance with relevant guidelines and regulations of Institutional Ethics Committee, Postgraduate Institute of Medical Education and Research, Chandigarh. Inclusion and Exclusion criteria is given in Supplementary methods. Baseline data of patients and control subjects is available in our earlier publication^16^.

### Renal artery ligation model of cardiac renal syndrome (RAL)

RAL was developed in adult wistar rats as described earlier^17^. Briefly, right renal artery of 20-week-old male rats (250–300 g; n =8) was ligated under anaesthesia, rats that underwent similar procedure without ligation served as sham-operated control group (n = 8). Animals were sacrificed 15 days after surgery and hearts were removed and either fixed in Karnovsky's fixative for histological study or immediately stored in liquid nitrogen for RNA and protein isolation. Cardiac hypertrophy was measured at the gross by heart weight (HW in mg) to body weight (BW in g) ratio. Expression of cardiac fibrosis (Col-1a, Col-4a CTGF, and FGF) and hypertrophy (ANP and β-MHC) genes at mRNA and protein levels was studied by qRT-PCR and western blotting respectively. Histopathological analysis (H&E staining and MT staining) was done in formalin fixed cardiac tissues to asses cardiac remodeling (hypertrophy and fibrosis). This study was approved by Institutional Animal Ethics Committee (IAEC), Postgraduate Institute of Medical Education and Research, Chandigarh and all experiments were done as per the guidelines and regulations of the committee.

### Modulation of Myocardin Gene

Myocardin (MYOCD) was transiently over-expressed in H9c2 cells or CFs by transfection with 2µg of pcDNA3-MYOCD-HA_v1, using lipofectamine^®^ 2000 transfection reagent (Invitrogen) according to the manufacturer’s instructions. For MYOCD cellular knockdown, H9c2 cells or CFs were transfected with 1µg/ml of MYOCD siRNA (Table S1 in supplementary data) which targets both the cardiac isoforms (Silencer^®^ Select pre-designed MYOCD siRNA; ID s140959; Ambion, USA). Cells transfected using scrambled sequence anti-miR (Ambion, USA) served as a negative control. Transfection efficiency was measured using Cy3 labelled negative siRNA by FACS Canto™ II Cell Analyser and by Olympus IX 73 Fluorescent microscopy.

### Cardiac specific silencing of MYOCD Gene

Cardiac MYOCD gene expression was inhibited in RAL rats using MYOCD specific siRNA, conjugated with a cardiac specific 20mer homing peptide sequence WLSEAGPVVTVRALRGTGSW encapsulated in stearic acid modified carboxy methyl chitosan as described earlier^18^, as per protocol shown in Supplementary Figure 1. Animals were sacrificed using sodium pantothenate as anaesthesia and performing cervical dislocation. Hearts were removed, examined grossly, and histopathological analysis (H&E staining and MT staining) was performed. RNA and protein was isolated from left ventricular tissue. Expression of hypertrophic (ANP and β-MHC) and pre-fibrotic genes (Col-1a, Col-3a, Col-4a CTGF, TGF β and FGF) was measured at mRNA and protein levels.

### Histological analysis

Tissues were fixed in 4% paraformaldehyde, embedded in paraffin, and sectioned at 4-μm thickness. Hematoxylin/eosin for cell size and morphology and Masson’s trichrome for collagen staining were performed using standard procedures. Nuclei were visualized using haematoxylin stain. Images were obtained by Olympus IX 73 microscope.

### MYOCD antibody specificity experiment

MYOCD antibody specificity was confirmed by immunostaining in cell lines which have high expression and sparse or null expression of MYOCD to ascertain that it is not binding to other family members of Myocardin like proteins and specific for MYOCD (Supplementary Figure 2). Imaging was done on Olympus FV1000 confocal microscopy.

### Angiotensin II (Ang II) induced cardiomyocyte hypertrophy

H9c2 rat cardiomyocytes were cultured in Dulbecco's Modified Eagle's Medium (DMEM), supplemented with 10% foetal bovine serum under 5% CO2 at 37°C. After 12 hrs of serum starvation, H9c2 cardiomyocytes were stimulated with 1µM angiotensin II (Ang-II) (Sigma Aldrich) or 7.5 ng/ml transforming growth factor-β (TGF-β) (Peprotech) for 24hrs, 48hrs and 72hrs. Expression of hypertrophic genes atrial natriuretic protein (ANP) was measured at mRNA and protein levels to confirm the development of hypertrophy.

### Ang II induced fibrosis in cardiac fibroblasts

Primary adult rat cardiac fibroblasts (CFs) were isolated from ventricular tissues of 6 to 8 week old male Wistar rats, (150-200gm) as previously described^19^. After 12hrs of serum starvation, CFs were stimulated with 1µM Ang II for 24hrs (these were optimal conditions for time and dose of Ang II as standardized in our laboratory for induction of fibrosis). Expression of various fibrotic genes (Col-1a, Col-3a, Col-4a CTGF, and FGF-b) was measured at mRNA levels. Institutional Animal Ethics Committee, Postgraduate Institute of Medical Education and Research, Chandigarh approved all animal experiments were performed in accordance with regulations specified by CPCSEA.

### Western Blotting

Cells and tissue protein lysate was prepared using RIPA lysis solution and 1X Protease inhibitor (Cat no.87785, Thermo Scientific), followed by sonication at 30% amplitude repeated for 3-4 times, 20secs each with 10secs interval. Protein concentration was estimated by quick start^TM^ Bradford dye reagent, 1X (Cat no. 500-0205, Bio-Rad). Laemmli Buffer was used for preparation of proteins. Protein lysate (50µg/well) was then separated on the basis of their molecular weight on SDS-PAGE Bio-Rad mini SDS-PAGE gel instrument. 50µg of protein sample along with 5µl of Page Ruler^TM^ pre-stained protein ladder (Cat no. 26617, Thermo Scientific), was loaded and gel was run initially at 50 volts until the tracking dye crosses the stacking gel, then at 100 volt till the dye come at the bottom of the gel. After SDS-PAGE, PVDF membrane was activated in 100% methanol and equilibrated in 1X transfer buffer along with SDS-PAGE gel and blotting sheets, at 4°C for 15 mins. Protein wet transfer was carried out at 30mA for 10hr in Mini-Protean Tetra System (Bio-rad). Ponceau staining was performed in order to check the protein transfer on PVDF membrane. PVDF membrane was then washed with 1X TBST buffer and kept in blocking solution (5% BSA in TBST buffer) for 2hrs on shaker at RT. After blocking, PVDF membrane was washed 3 times in TBST buffer (1X) for 10mins each and was then incubated in primary antibody (List of the antibodies used in the study with the working dilution, source and catalogue number are provided as Table S2 in Supplementary data) for 4°C overnight. PVDF membrane was then washed 3 times with 1X TBST buffer for 10mins each and then incubated with HRP labelled secondary antibody for 40 mins at RT. PVDF membrane was then washed 3times with 1X TBST buffer for 10mins each and target protein reactive with primary antibodies was detected. After incubation with the appropriate secondary antibodies, blots were visualized using the EZ-ECL detection system (Cat no.20-500-120A, Biological Industries) and the FluorChem M luminescent image analyzer (Protein Simple, Biotechne brand, USA). Densitometry analysis of band intensity was performed using Image J software (NIH, USA).

### Statistical Analysis

The statistical analysis was carried out using Statistical Package for Social Sciences (SPSS Inc., Chicago, IL, and version 21.0 for Windows)/Graph Pad Prism 5.0. All quantitative variables were estimated using mean and measures of dispersion (standard deviation and standard error). Normality of data was checked by measures of skewness and Kolmogorov Smirnov tests of normality. Expression of genes were compared using Mann Whitney or Student’s t-test depending on the normality of data. One way/Two way ANOVA with appropriate post-hoc tests were performed when multiple comparisons were done within various experimental groups. All statistical tests were two-sided and performed at a significance level of p≤0.05*, p≤0.01** and ***p≤0.001.

## Results

### Cardiac MYOCD expression level is significantly increased in cardiorenal syndrome patients

We analyzed the mRNA and protein expression levels of MYOCD in endomyocardial biopsies of DCM patients with and without renal failure. The mRNA and protein expression levels of MYOCD was increased in endomyocardial biopsies obtained from DCM patients compared to healthy controls. However, the increase is more pronounced in DCM patients with renal failure compared to DCM alone patients. (Figure 1a & b).

**Fig 1.**
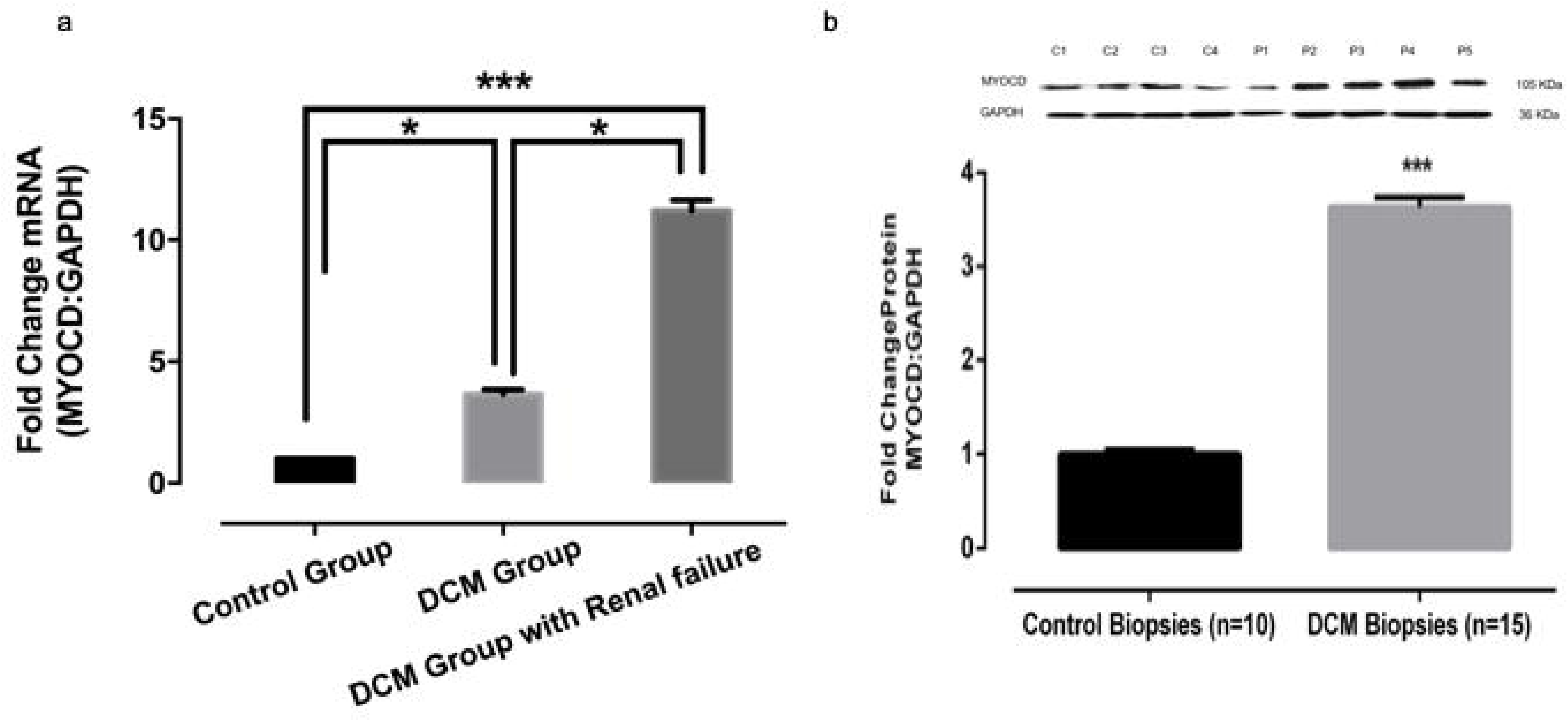
Myocardin (MYOCD) mRNA and protein expression in myocardial tissues of DCM patients with and without renal failure and controls. **a)** Quantitative mRNA expression of MYOCD in DCM cardiac tissues (n=30) with and without renal failure and control group (n=15) as determined by qRT-PCR using Taqman probe chemistry. **b)** Representative western blot for MYOCD and GAPDH proteins in control (C1–C4) and DCM patients (P1–P5) in LV myocardium samples. Semi-Quantitative protein expression of MYOCD in DCM patients and controls as estimated by software Image J. Normalization of total RNA and protein was done using GAPDH as an internal control. Data given is Mean ± SEM; *p<0.05 Control vs DCM; ***p<0.001 Control vs DCM

### MYOCD activation induces hypertrophic and fibroblast associated genes

To know the functional relevance of increase of MYOCD in cardiorenal patients, we treated cardiomyocytes cells (H9c2) and fibroblasts with Ang II (a well-known modulator of cardiorenal disease). The treatment showed increased mRNA and protein expression of MYOCD in both cell types (Supplementary Figure 3a, 3b & 3c), along with hypertrophy markers including Atrial Natriuretic factor (ANP), Beta-Myosin heavy chain (β-MHC) and fibrotic genes at mRNA levels such as Collagen (Col) 1a, Col 3a, Col 4a, Transforming growth factor-β (TGF-β), Connective tissue growth factor (CTGF) and Fibroblast growth factor (FGF)-β respectively (Figure 2a, 2b & 2c). Pre-treatment with losartan (an Ang II type I receptor antagonist) reversed these effects by increasing the levels of MYOCD, ANP and β-MHC mRNA levels in cardiomyocytes, whereas, treatment with known Ang II analogue had no significant effect on MYOCD, ANP and β-MHC expression in these cells (Supplementary Figure 3d).

**Fig 2.**
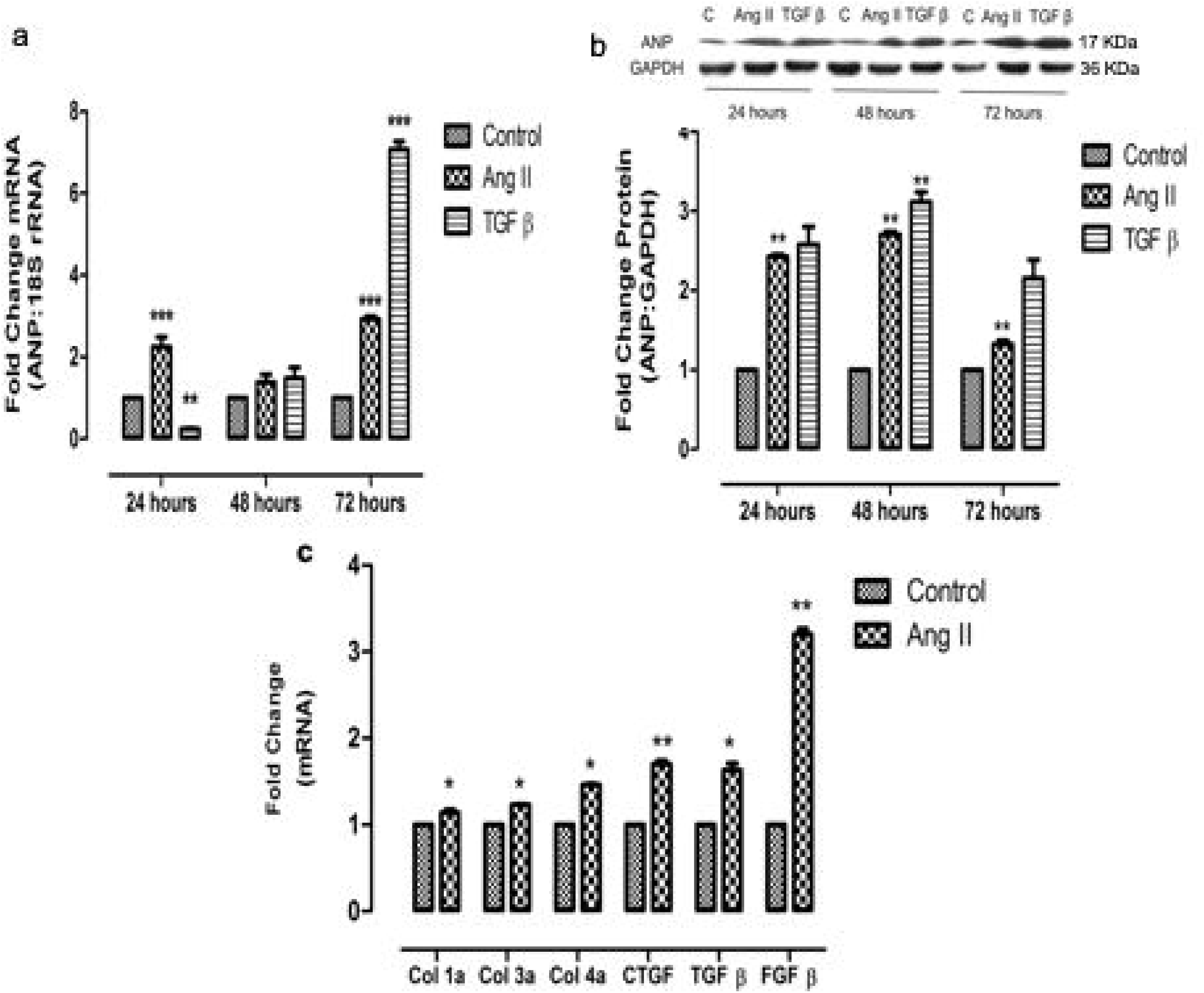
Effect of MYOCD overexpression on Ang II induced expression of hypertrophy and fibrosis genes in H9c2 cardiomyocytes and fibroblasts respectively. **a)** Quantitative mRNA expression of Atrial Natriuretic factor (ANP) in Ang-II (1µM) and TGF β (7.5 ng/ml) treated H9c2 cells as determined by qRT-PCR. **b)** Representative western blot for ANP and GAPDH proteins in Ang-II and TGF-β treated H9c2 cells. **c)** Quantitative mRNA expression of Collagen (Col) 1a, Col3a, Col4a, CTGF, TGF-β and FGF-β (fibrotic marker gene) in Ang II treated cardiac fibroblasts as determined by qRT-PCR. Normalization of total RNA and protein was done by using GAPDH as an internal control. Data given is Mean ± SEM of triplicates; *p<0.05; **p<0.01; ***p<0.001 Control vs Ang II and Control vs TGF β.

### *In vitro* silencing of MYOCD inhibited Ang II induced cardiomyocyte hypertrophy and fibrosis

To examine the effect of MYOCD silencing on cardiomyocyte hypertrophy, Ang II treated H9c2 cells were transfected with MYOCD siRNA. siRNA transfection significantly reduced the MYOCD levels (Supplementary Figure 4a & 4b), leading to the reduction of ANP and β-MHC levels (Figure 3a & 3b).

**Fig 3.**
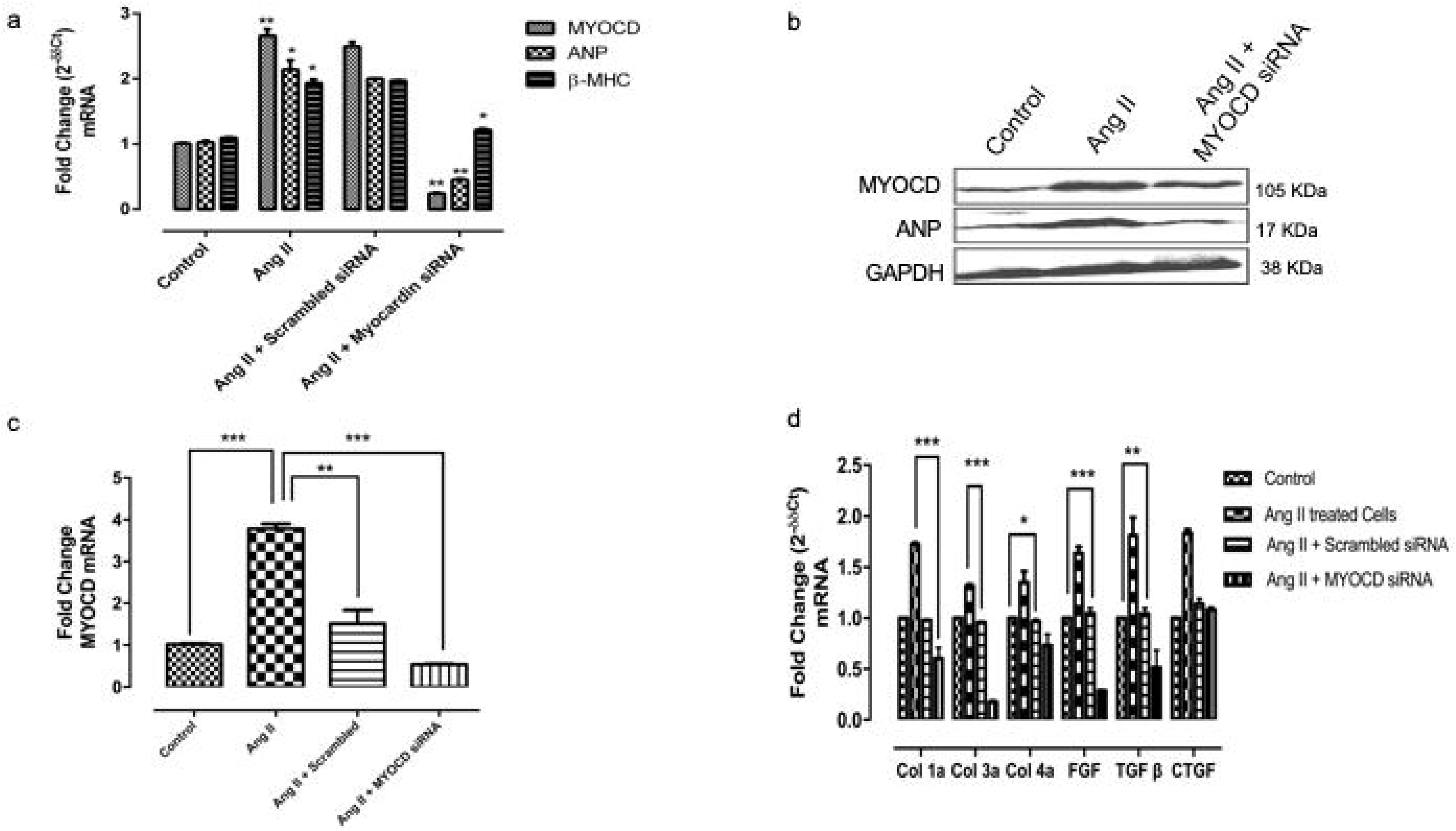
MYOCD siRNA treatment on expression of hypertrophic genes and fibrotic genes in H9c2 and cardiac fibroblast respectively. **a)** mRNA expression of ANP, Beta myosin heavy chain (β-MHC) and MYOCD after treatment with MYOCD siRNA in Ang II treated H9c2 cells **b)** ANP and MYOCD protein expression after treatment with MYOCD siRNA in Ang II treated H9c2 cells **c)** mRNA expression of MYOCD expression after treatment with MYOCD siRNA in cardiac fibroblast cells **d)** mRNA expression of fibrotic gene (Col1a, Col3a, Col4a, CTGF, TGF β and FGF β) after treatment with MYOCD siRNA in Ang-II treated fibroblast cells. Normalization of total RNA and protein was done by using GAPDH as an internal control. Data given is Mean ± SEM of triplicates; *p<0.05; **p<0.01; ***p<0.001Control vs Ang II and Ang II vs Ang II + MYOCD siRNA

Also, treatment of MYOCD siRNA significantly decreased MYOCD mRNA expression levels in the cardiac fibroblasts (Figure 3c), whereas scrambled siRNA also showed some effect on MYOCD expression in these cells but the changes found in myocd siRNA treated cells was much significant. MYOCD siRNA treated cells resulted in a significant decrease in mRNA expression of Col 1a, Col 3a, Col 4a,TGF-β and FGF-β genes in Ang II treated cardiac fibroblasts (Figure 3d). We found changes in CTGF expression both with Ang II treatment and myocd siRNA but those changes were non-significant.

### Cardiac-specific inhibition of MYOCD gene in a cardiorenal rat model improves left ventricular functions

To know whether MYOCD inhibition in a cardiorenal rat model can rescue the above observed *in vitro* phenotype, we performed renal artery ligation (RAL) in rats. This model has been shown to an excellent model for cardiac-renal heart failure^20^. RAL in rats resulted in cardiac hypertrophy and systolic dysfunction (Figure 4a & 4b; and Table 1).

**Fig 4.**
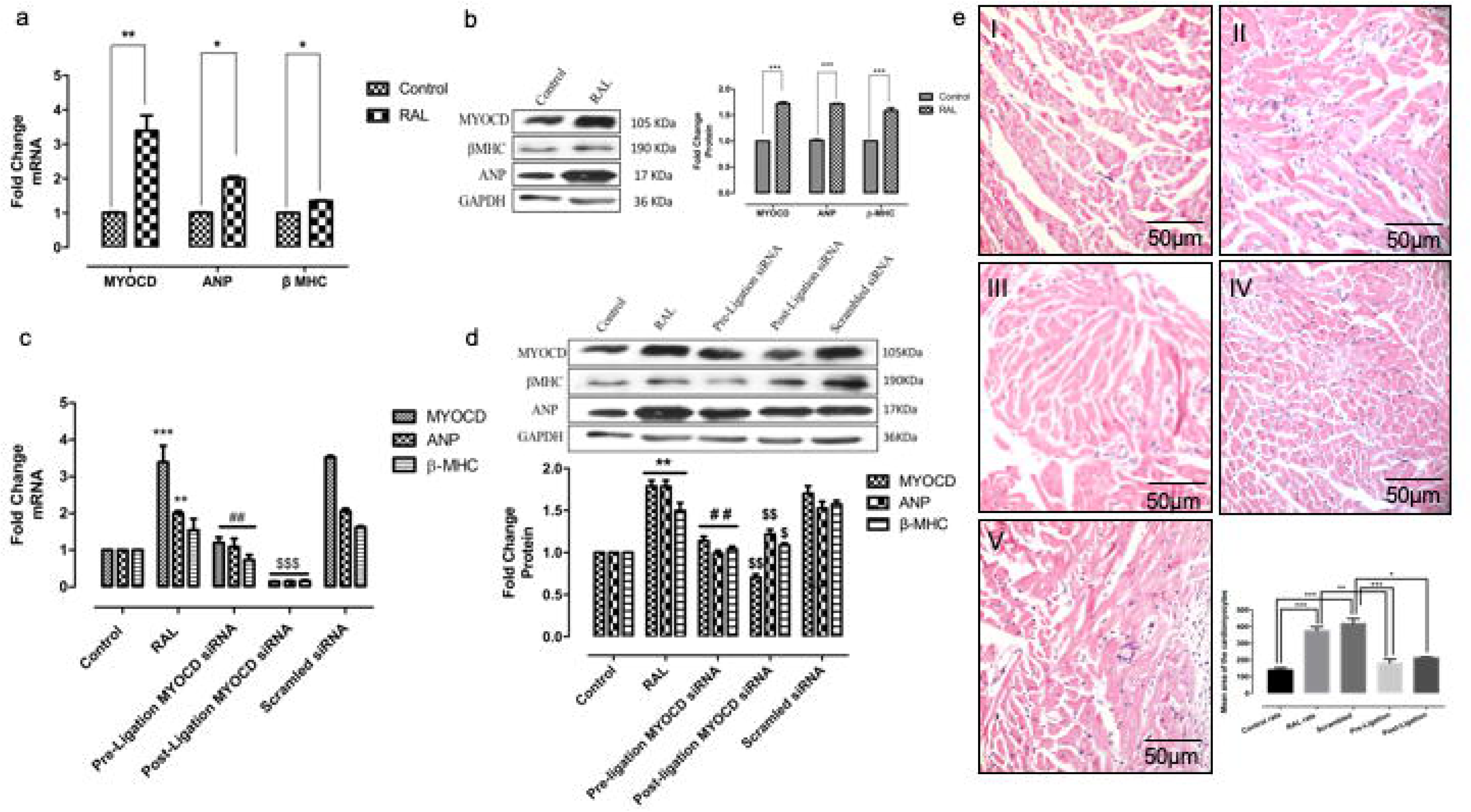
Effect of MYOCD siRNA treatment on expression of hypertrophic genes (ANP and β-MHC) and on cardiomyocyte size in RAL rats. **a)** MYOCD, ANP and β-MHC mRNA expression in RAL rats (n=8) **b)** Representative western blot for MYOCD, ANP and GAPDH proteins in RAL rats. **c)** mRNA expression of MYOCD, ANP and β-MHC in MYOCD siRNA treated RAL rats **d)** Representative western blot for MYOCD, ANP and GAPDH proteins in MYOCD siRNA treated RAL rats (before and after ligation). **e)** Haematoxylin and Eosin (H&E) stained Photomicrographs (10X) of rat hearts: I) Control group, II) RAL group III) Scrambled treated RAL group IV) MYOCD siRNA treated (before ligation) RAL Group V) MYOCD siRNA treated (after ligation) RAL group. Bar diagram shows mean of cardiomyocyte area (n=50) as measured by Image J analyser. Normalization of total RNA and protein was done using GAPDH as an internal control. Data given is Mean ± SEM; **p<0.01; ***p<0.001 for Control vs RAL; ^##^p<0.01 for RAL vs RAL treated with MYOCD siRNA before ligation (Pre-ligation); ^$^p<0.05, ^$$^p<0.01 ^$$$^p<0.001 for RAL vs RAL treated with MYOCD siRNA after ligation (Post-Ligation)

**Table 1:** Echocardiographic parameters and heart weight (HW) to body weight (BW) ratio in RAL model treated with myocd siRNA

| Treatment Groups | LVDD (mm) | FS (%) | LVEF (%) | HW/BW ratio |
| --- | --- | --- | --- | --- |
| Group I (Sham Controls) | 4.34 ± 0.20 | 62.08 ± 1.18 | 78 ± 8.8 | 3.2 ± 0.5 |
| Group II (Ligated) | 5.83 ± 0.38* | 34.6 ± 3.8* | 55 ± 6.7* | 4.6 ± 0.7* |
| Group III (Ligated + post-myocd siRNA) | 4.95 ± 0.67<sup>#</sup> | 56.8 ± 1.97<sup>#</sup> | 72 ± 5.9<sup>#</sup> | 3.95 ± 0.3<sup>#</sup> |
| Group IV (Ligated+ pre-myocd siRNA) | 5.05 ± 0.45<sup>\$</sup> | 50.6 ± 2.85<sup>\$</sup> | 61 ± 7.2<sup>\$</sup> | 3.7 ± 0.4<sup>\$</sup> |
| Group V (Ligated+ scrambled siRNA) | 5.6 ± 0.4 | 38 ± 2.4 | 52 ± 4.4 | 4.7 ± 0.5 |

LVDD: Left Ventricular Diastolic Diameter; FS: Fraction Shortening; LVEF: Left Ventricular ejection fraction. \*p<0.05 Group I vs Group II; <sup>#</sup>p<0.05 Group II vs Group III; <sup>\$</sup>p<0.05 Group II vs Group IV

Next, we injected the MYOCD siRNA conjugated with homing peptide through tail veins of the RAL rats as described earlier to be cardiac-specific and having no bystander effects^18^. To study the effect of heart-specific MYOCD silencing on cardiac function, echocardiography and HW/BW ratio were measured in MYOCD siRNA treated RAL rats 15 days post ligation. HW/BW ratio was significantly reduced in MYOCD siRNA treated RAL rats (Table 1). A significant decrease in LVEDD and increase in LVEF was also seen in these rats (Table 1) indicating improvement in left ventricular function by MYOCD inhibition.

### MYOCD inhibition in the RAL model attenuated cardiac hypertrophy and fibrosis

Ablation of MYOCD gene with MYOCD siRNA conjugated with homing peptide in RAL rats resulted in significant reduction in cardiac MYOCD expression (Figure 4c & d), whereas scrambled siRNA did not affect the MYOCD expression. Also, a profound reduction in the expression of hypertrophy markers, ANP and β-MHC (Figure 4c & d) and a decrease in cardiomyocyte size in MYOCD siRNA treated RAL rats compared to untreated group were observed (Figure 4e III & IV). There was no evidence of inflammation or coronary occlusion in these rats, suggesting absence of ischemic lesions (Fig. 4e).

This model showed cardiac fibrosis along with hypertrophy (Figure 5a & b). Furthermore, MYOCD siRNA treatment, resulted in a significant reduction in cardiac fibrotic gene expressions including Col 1a, Col 3a, Col 4a, TGF β, CTGF and FGF β compared to untreated RAL rats (Figure 5 c & d). Histopathological examination showed decreased collagen deposition in the siRNA treated RAL cardiac tissues (Figure 5e). Collectively, these data suggested that the MYOCD inhibition can reverse the heart failure in RAL model.

**Fig 5.**
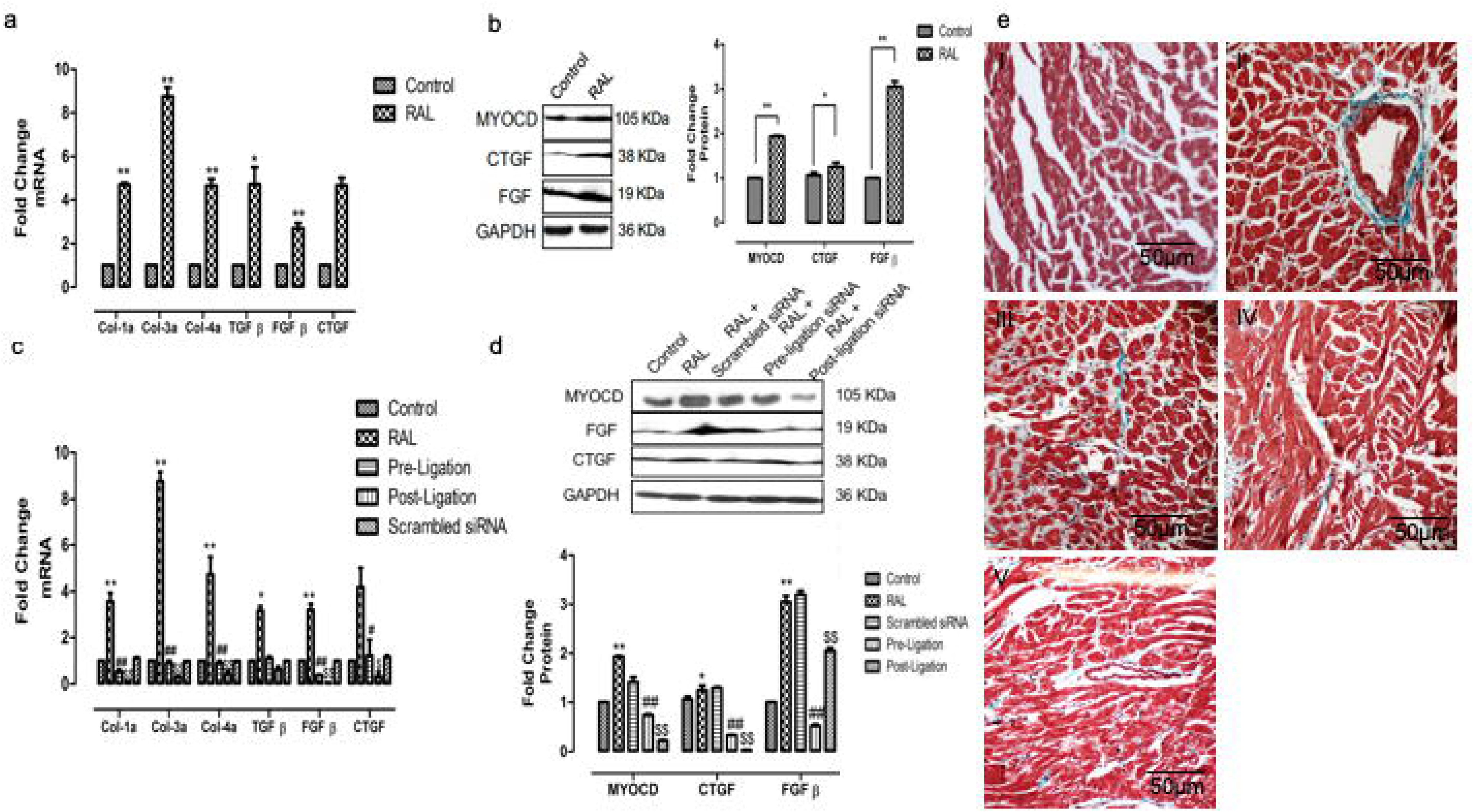
Effect of MYOCD siRNA treatment on expression of fibrotic genes (Col 1a, Col 3a, Col 4a, TGF β, CTGF and FGF β) in RAL rats (n=8) **a)** mRNA expression of Col 1a, Col 3a, Col 4a, TGF β, CTGF and FGF β in RAL rats. **b)** Representative western blot for CTGF, FGF and GAPDH proteins in RAL rats. **c)** mRNA expression of Col 1a, Col 3a, Col 4a, TGF β, CTGF and FGF β in MYOCD siRNA treated RAL rats (before and after ligation). **d)** Representative western blot for CTGF, FGF and GAPDH proteins in MYOCD siRNA treated RAL rats (before and after ligation). **e)** Photomicrographs (10X) of rat heart tissues with Masson’s Trichrome staining which is collagen specific and evaluates fibrosis: I) control, II) RAL rats with extensive collagen deposition III) Scrambled treated RAL group III) MYOCD siRNA (before ligation) treated RAL rats show decreased collagen deposition IV) MYOCD siRNA (after ligation) treated RAL rats show decreased collagen deposition. Normalization of RNA and protein was done using GAPDH as an internal control. Data given is Mean ± SEM; *p<0.05; **p<0.01 for Control vs RAL; ^#^p<0.05; ^##^p<0.01 for RAL vs RAL treated with MYOCD siRNA before ligation; ^$^p<0.05; ^$$^p<0.01 for RAL vs RAL treated with MYOCD siRNA after ligation.

## Discussion

MYOCD role have been reported in various animal models of HF and in end-stage heart failure patients^14,15^. However, its role in cardiorenal patients remains unexplored. We observed a significant upregulation of cardiac MYOCD in endomyocardial biopsies from cardiorenal patients. This is the first observation on the MYOCD expression levels in the cardiorenal failure patients.

Cardiorenal syndrome patients display a significant amount of circulating renin with Ang II production, leading to arteriolar constriction and high venous pressure. Ang II along with aldosterone release result in systemic pressure through increased sodium re-absorption in the distal nephrons. These ultimately worsen the patient cardiac functions^21^. Thus, consistent activation of RAAS contributes to poor prognosis of the cardiorenal patients. RAAS activation is shown to be associated with upregulation of MYOCD related transcription factors leading to increased transforming growth factor-β (TGF-β) and fibronectin^11,22^. In the present study, direct inhibition of MYOCD in Ang II treated cardiomyocytes and fibroblasts resulted in decreased expression of hypertrophic and pro-fibrotic markers including TGF-β. These data suggest RAAS activation can be mitigated by MYOCD inhibition.

Torrado et al. showed that moderate inhibition of MYOCD using shRNA in doxorubicin-induced HF (DHF) model could partly restore diastolic dysfunction and decreased the DHF animal mortality^23^. However, such an effect remains to be explored in a cardiorenal model. We explored whether heart-specific MYOCD targeting could ameliorate cardiac remodeling and improve cardiac function in a rat renal artery ligation (RAL) model. RAL is a well-established model for studying the cardiac remodeling in renal dysfunction associated heart failure^20^.

Notably, we explored if MYOCD silencing pre-or post-ligation by siRNA could have effective in a RAL rat model showing increased expression of cardiac MYOCD with extensive cardiac hypertrophy and fibrosis. A cardiac specific homing peptide conjugated MYOCD siRNA to specifically inhibit cardiac MYOCD without any specific bystander effects was employed^18^. We observed that this cardiac specific MYOCD silencing resulted in attenuation of cardiomyocyte hypertrophy and cardiac fibrosis when given at either both at pre-ligation or post-ligation stages.

Further, we compared whether siRNA inhibition was more effective at pre- or post-ligation to ascertain the efficacy of MYOCD inhibition on cardiac hypertrophy and fibrosis under these two conditions. In pre-ligation treatment, MYOCD siRNA was introduced before the initiation of cardiac remodeling process, and in post-ligation treatment, MYOCD siRNA was given after the initiation of cardiac remodeling. Both the treatment groups showed significant reduction in the cardiac remodeling and improvement of cardiac function, but post-ligation treatment showed a greater reduction. These results suggest that MYOCD inhibition is more effective at post ligation stage, probably as MYOCD upregulation is maximal at this stage.

Our data suggest cardiac specific MYOCD silencing in cardiorenal rats model (RAL) resulted in the significant reduction in expression of hypertrophy markers, ANP and β-MHC and decrease in cardiomyocyte size. Further, the histopathological analysis also revealed reduction in cardiomyocyte size will be beneficial in ameliorating the process of cardiac hypertrophy. A significant decrease was observed in the cardiac expression of Col 1a, Col 3a, Col 4a, TGF-β, CTGF and FGF-β genes in MYOCD siRNA treated RAL rats. Histopathological examination corroborated the beneficial effect of MYOCD siRNA, as myocardial collagen deposition decreased in siRNA treated rats, suggesting the potential in controlling the cardiac fibrosis. MYOCD silencing was done in a controlled manner so there are less chances of bystander effects on the normal physiology of the myocytes and fibroblasts. Cardiomyocytes were found to have normal morphology and functional capacity after the MYOCD siRNA treatment.

Our data document for the first time, the upregulation of cardiac MYOCD in cardio-renal patients and cardiac-specific MYOCD silencing in a cardiorenal model has an integrated beneficiary effect on cardiac hypertrophy, fibrosis and function. Our data also strengthen the claims that cardiac targeted MYOCD inhibition might be a potential therapeutic target in reversing cardiac remodeling in the cardiorenal syndrome.

## Acknowledgement and Funding Sources

We would like to acknowledge Prof. Shyam K. Singh, PGIMER, Chandigarh for providing us with the control tissues. We are also thankful to Prof. Joseph A Miano, Rochester Institute, USA for providing MYOCD cardiac isoform plasmids as a kind gift. Anupam Mittal was recipient of DS Kothari Postdoctoral Fellowship, UGC and National Postdoctoral Fellowship by DST SERB. PSD is funded by American Heart Association-Scientist Development Grant and Welcome Trust-Intermediate Fellowship. While this manuscript is in review, Lyu et al., reported a size variability for MYOCD protein. Further studies using the antibodies reported by Lyu et al will further confirm our findings^24^.

## Contribution Statement

Anupam Mittal, Santanu Rana, Akhilesh Kumar and Satish K Raut were involved in acquisition of data. Anupam Mittal, Madhu Khullar and Perundurai Dhandapany was involved in analysis and interpretation of data. Anupam Mittal, Madhu Khullar, and Perundurai Dhandapany were involved in drafting of the manuscript. Ajay Bahl, Uma Nahar Saikia, Rishikesh Prasd and Sagartirtha Sarkar were involved in technical support for patient data. Rajni Sharma was involved in statistical analysis and analysis and interpretation of data.

