## Supplementary Data and information for "Myocardin ablation in a cardiac-renal rat model"

**Short/Running Title: Myocardin ablation in a cardiac-renal rat model**

| **Table S1: Sequences of SYBR Green Chemistry Primers and siRNAs** | | | | |
| --- | --- | --- | --- | --- |
| **Genes** | **Forward Primer** | **Reverse Primer** | **Product Size (bp)** | **T_m_**  **(°C)** |
| **Rno-Myocardin** | 5’AGATCCCGGC  CGCTTCCCTTT’3 | 5’GGTCGGGAAT  CTGGGAGAGG’3 |  | 58 |
| **Rno-CTGF** | 5’GAGTCGTCTC  TGCATGGTCA‘3 | 5’CCACAGAACT  TAGCCCGGTA‘3 | 156 | 58 |
| **Rno-FGF-β** | 5’CGGTACCTGG  CTATGAAGGA‘3 | 5’CCGTTTTGGA  TCCGAGTTTA‘3 | 178 | 58 |
| **Rno-TGF-β** | 5’TGCTTCAGCTC  CACAGAGAA‘3 | 5’TGGTTGTAGA  GGGCAAGGAC‘3 | 182 | 58 |
| **Rno-Col-1a** | 5’TGCTGCCTTT  TCTGTTCCTT‘3 | 5’AAGGTGCTGG  GTAGGGAAGT‘3 | 179 | 58 |
| **Rno-Col-3a** | 5’CATCTTTTCCA  GGAGGTCCA’3 | 5’GTCCACGAGGT  GACAAAGGT‘3 | 189 | 58 |
| **Rno-Col-4a** | 5’GCCAAGTGTG  CATGAGAAGA‘3 | 5’AGCGGGGTGT  GTTAGTTACG‘3 | 202 | 58 |
| **Rno-ANP** | 5’ATTTCAAGAACC  TGCTAGACC’3 | 5’TTTTCAAGAG  GGCAGATCTAT’3 | 222 | 58 |
| **Rno-β-MHC** | 5’CCTCGCAATAT  CAAGGGAAA’3 | 5’TACAGGTGCAT  CAGCTCCAG’3 | 198 | 58 |
| **Hsa/Rno-GAPDH** | 5’ACAGCAACAG  GGTGGTGGAC‘3 | 5’TTTGAGGGTG  CAGCGAACTT‘3 | 252 | 58 |
| **Rno-Myocardin siRNA 1** | 5’GGUCAAACCCA  UGUACUCUTT‘3 | 5’AGAGUACAUGGG  UUUGACCTG‘3 |  |  |

Atrial Natriuretic factor (ANP), Beta-Myosin heavy chain (β-MHC), Collagen (Col) 1a, Col 3a, Col 4a, Transforming growth factor-β (TGF-β), Connective tissue growth factor (CTGF) and Fibroblast growth factor (FGF)- β

| **Table S2: Source and Working Concentration information of antibodies** | | | |
| --- | --- | --- | --- |
| **S.No.** | **Antibody** | **Protein MW (kDa)** | **Dilution Factor**  **(DF)** |
| **1** | **Myocardin (Sigma; #SAB4200539)** | 105 kDa | 1:1000 |
| **2** | **ANP (Thermo Pierce; sc-18811)** | 17 kDa | 1:750 |
| **3** | **β-MHC (Santacruz; sc-168678)** | 190 kDa | 1:1000 |
| **4** | **GAPDH (Sigma; G9545)** | 36 kDa | 1:1000 |
| **5** | **CTGF (Santacruz; sc-365970)** | 38 kDa | 1:700 |
| **6** | **FGF (Santacruz; sc-1390)** | 19 kDa | 1:1000 |
| **7** | **Anti-mouse IgG-HRP**  **(Santacruz; A9919)** |  | 1:10000 |
| **8** | **Anti-goat IgG-HRP**  **(Santacruz; A5420)** |  | 1:10000 |
| **9** | **Mouse anti-rabbit IgG-HRP**  **(Santacruz; sc-2357)** |  | 1:10000 |

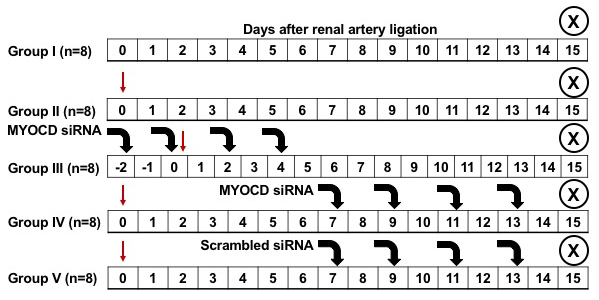
**Supplementary Figure 1: Experimental Design of siRNA treatment in RAL group:** 20 weeks old wistar rats were randomized in five groups, assigned to undergo ligation of right renal artery and SHAM procedure. Group I: SHAM operated rats (Control group), Group II: Ligated rats (RAL group), Group III: Myocardin (MYOCD) siRNA introduced in ligated group after ligation and initiation of cardiac remodelling (Post-Ligation group), Group IV: MYOCD siRNA introduced in ligated group before ligation and initiation of cardiac remodelling (Pre-Ligation group). Group V: Scrambled siRNA introduced in ligated group. Red arrow shows the day of ligation. Black coloured bold arrow shows days on which MYOCD siRNA/Scrambled negative siRNA was introduced and Encircled cross shows day of sacrificing rats.

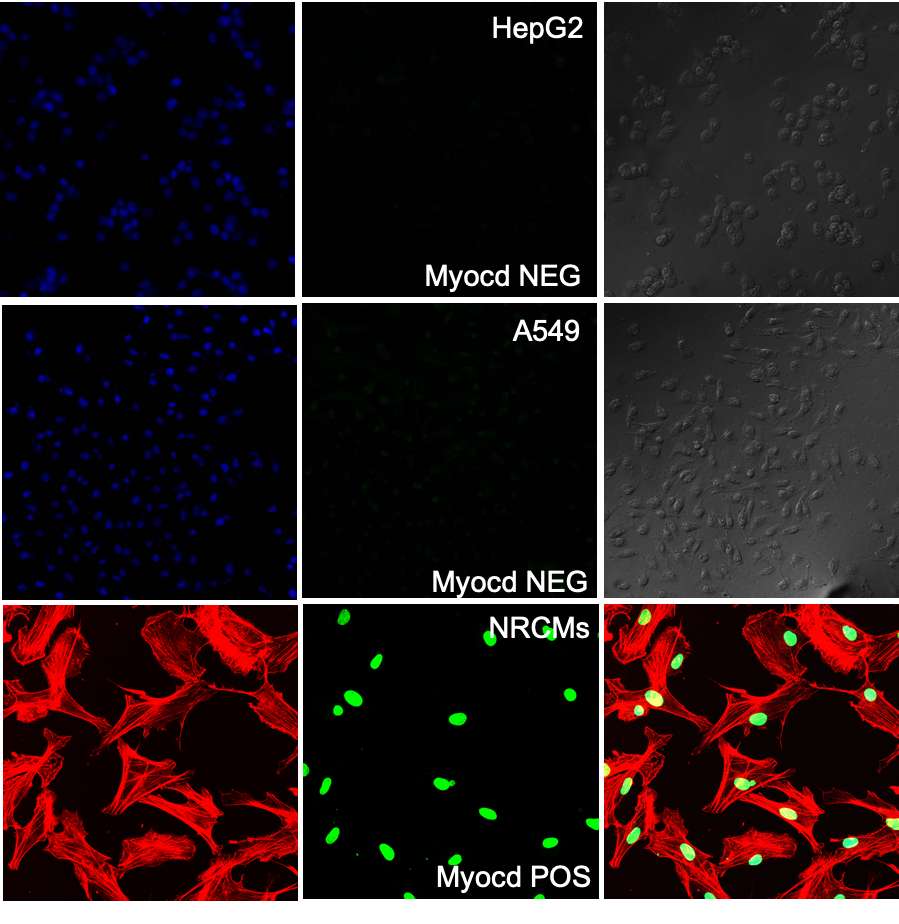

**Supplementary Figure 2: Myocardin (MYOCD) immunostaining in various cell lines to confirm the specificity of the antibody.** Upper Panel: Immunocytochemistry done in HepG2 hepatocarcinoma cells (Anti-MYOCD dilution: 1:200) shows negative expression of MYOCD. Middle Panel: A549 cells stained with same dilution of MYOCD shows very faint (nearly negatively) stained cells. Lower Panel: Neonatal rat cardiomyocytes (NRVCs) stained with 1:200 anti-MYOCD show good MYOCD expression. MYOCD antibody was used even at lower dilution of 1:100 in HepG2 and A549 but showed similar results (data not shown).

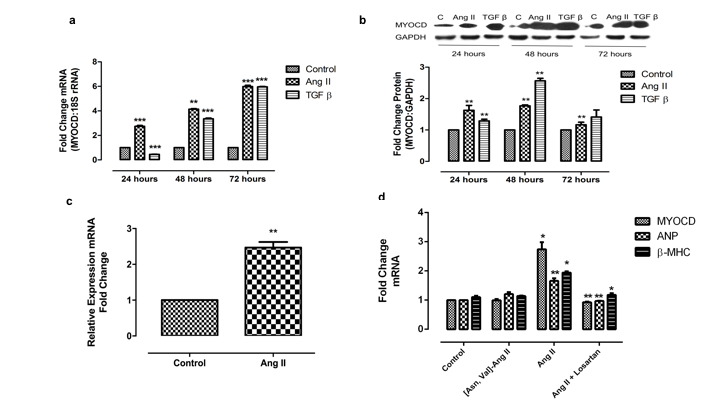

**Supplementary Figure 3: Myocardin (MYOCD) expression in Ang-II and TGF β treated H9c2 cardiomyoctes and cardiac fibroblast cells.** H9c2 cardiomyocytes and cardiac fibroblast were incubated with 1µM of Ang II for 24 hours to induce hypertrophy and fibrosis **a)** MYOCD mRNA expression was determined by the qRT-PCR in H9c2 cardiomyocytes and **b)** Representative blot showing the expression of MYOCD in time dependent manner in H9c2 cardiomyocytes **c)** MYOCD mRNA determination in Ang-II treated cardiac fibroblasts d) Angiotensin II type I receptor antagonist, Losartan treatment in Ang II treated H9c2 confirmed that increase in MYOCD and hypertrophy genes (ANP and β-MHC) is mediated by Ang II. GAPDH was used for normalization of the qRT-PCR and protein expression analysis. Data given is Mean ± SEM. *p<0.05, **p<0.01, ***p<0.001 Control vs Ang II; Control vs TGF β.

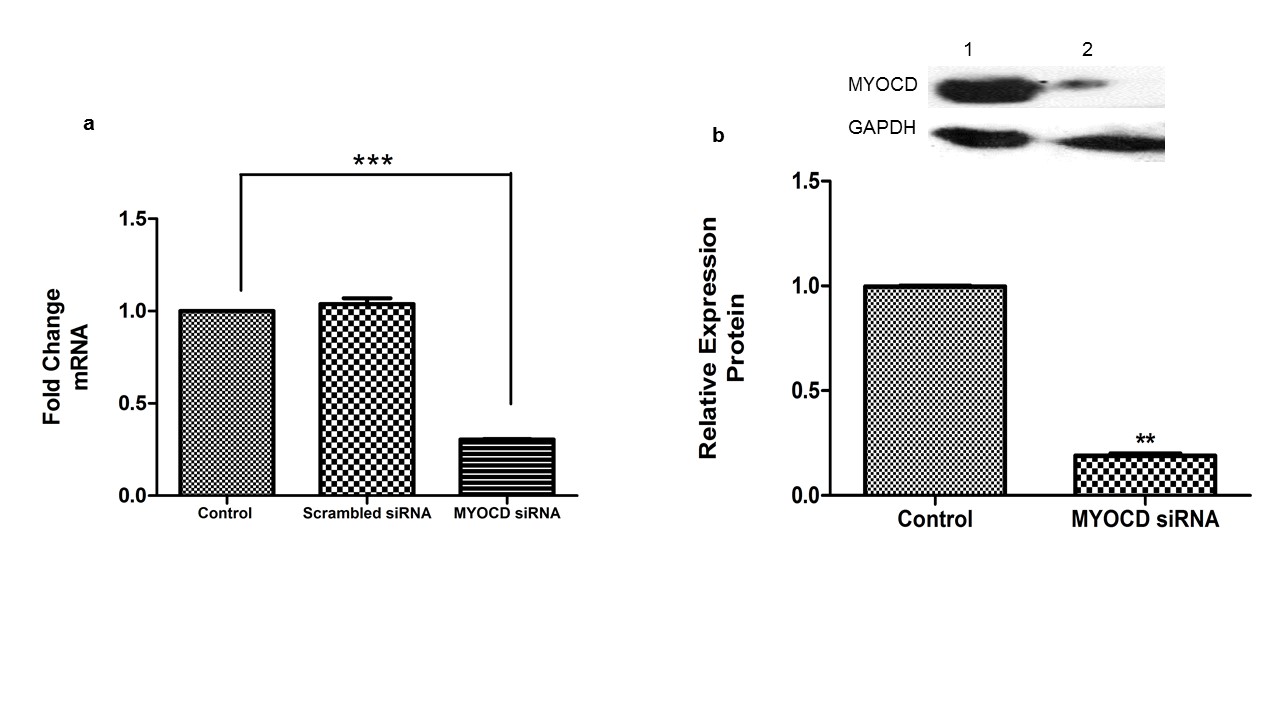
**Supplementary Figure 4: Expression of Myocardin (MYOCD) after transfection with MYOCD specific siRNA into the cardiomyocytes. a)** MYOCD mRNA expression after transfection with MYOCD siRNA. **b)** Representative western blot and quantitative results of the MYOCD and GAPDH protein expression. Lane1: Cells with only vector, Lane 2: Cells transfected with MYOCD siRNA. Normalization was done using GAPDH as internal control. Data shown are results from three independent experiments run in triplicate. Data given is Mean ± SEM; **p<0.01, ***p<0.001 Control vs MYOCD siRNA.

**Supplementary Information**

Figure 1: Western Blot showing MYOCD expression in all control (n=10) and DCM patient biopsies (n=15) other than which has been shown as representative images in the main figure 1b

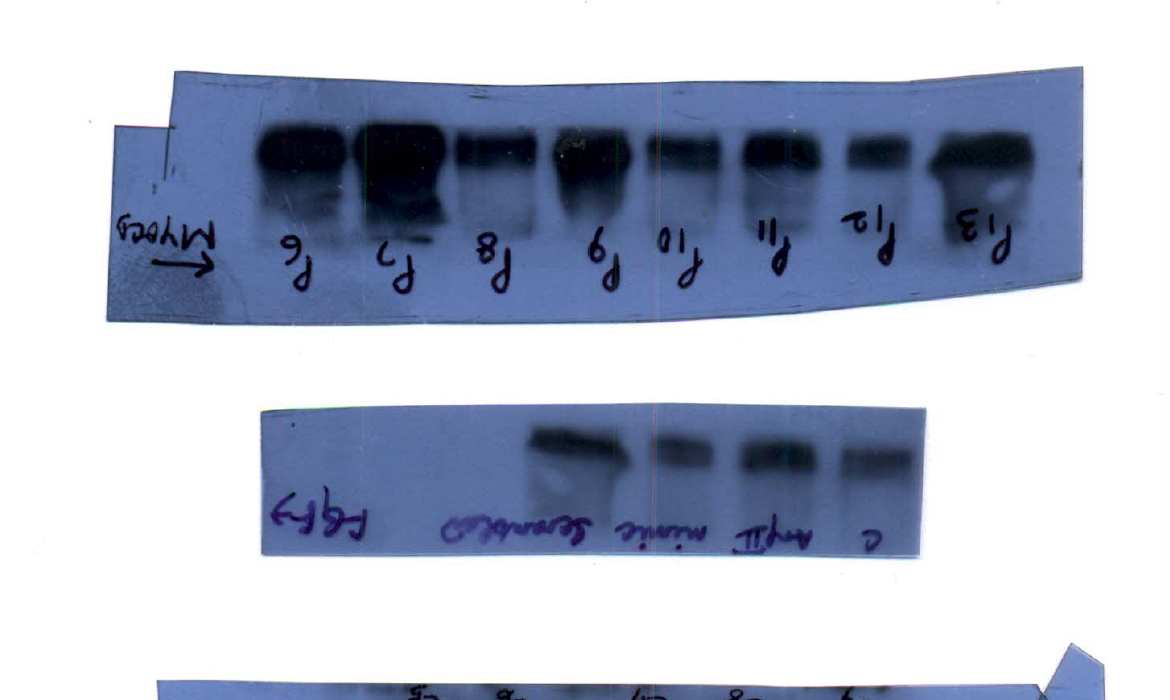

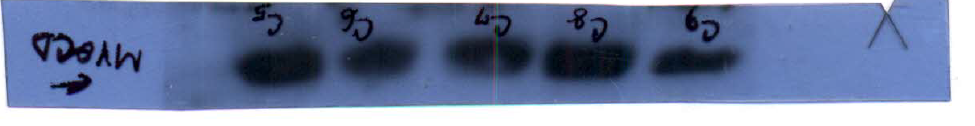

Figure 2: Raw western blots showing ANP expression after Ang II and TGF-β in time dependent manner in H9c2 cells (Sequence from left to right 1. Control 24 hrs, 2. Ang II 24 hrs, 3. TGF- β 24 hrs, 4. Control 48 hrs, 5. Ang II 48 hrs, 6. TGF- β 48 hrs, 7. Control 72 hrs, 8. Ang II 72 hrs, 3. TGF- β 72 hrs)

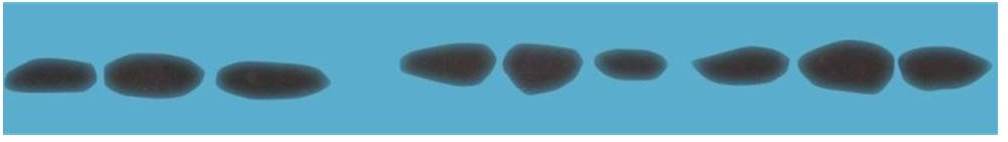

GAPDH

ANP

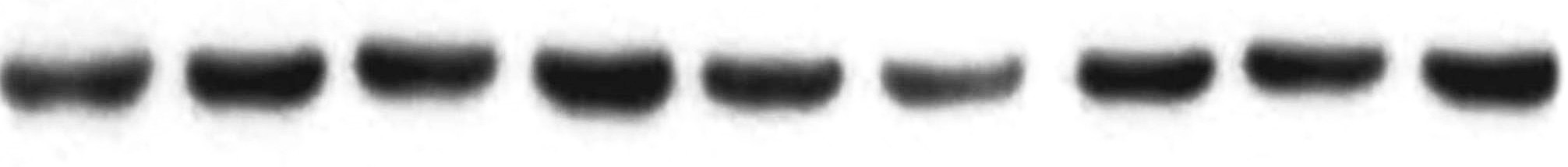

Figure 3: Unprocessed western blot showing ANP and MYOCD protein expression after treatment with MYOCD siRNA in Ang II treated H9c2 cells (Sequence from Left to right: 1. Control H9c2, 2. Ang II, 3. Ang II + MYOCD siRNA1, 4. Ang II + MYOCD siRNA 2). siRNA2 data is not shown in Final figure

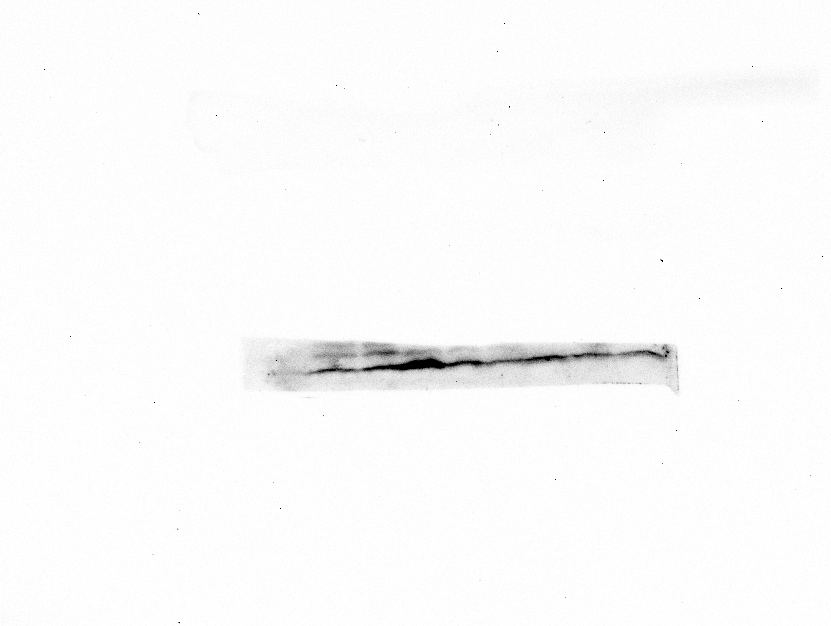

ANP

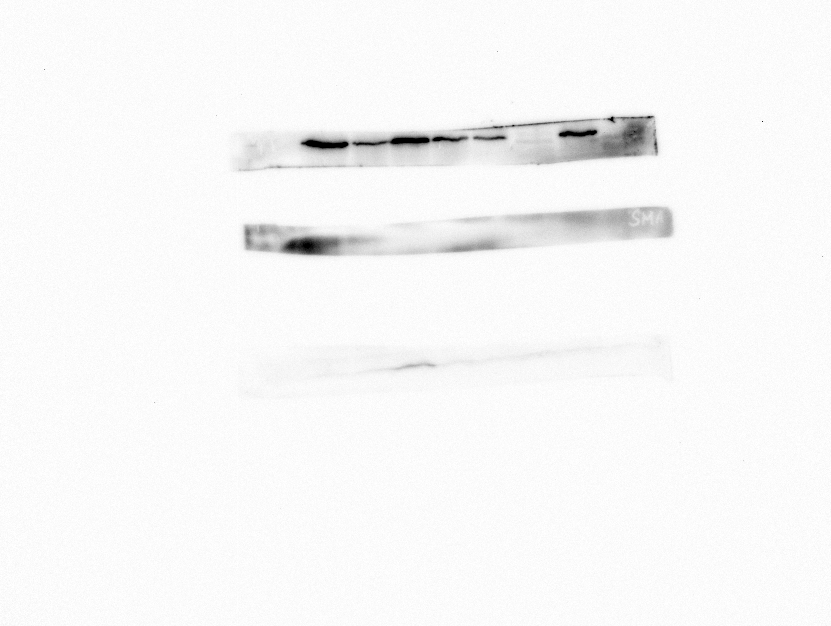

MYOCD

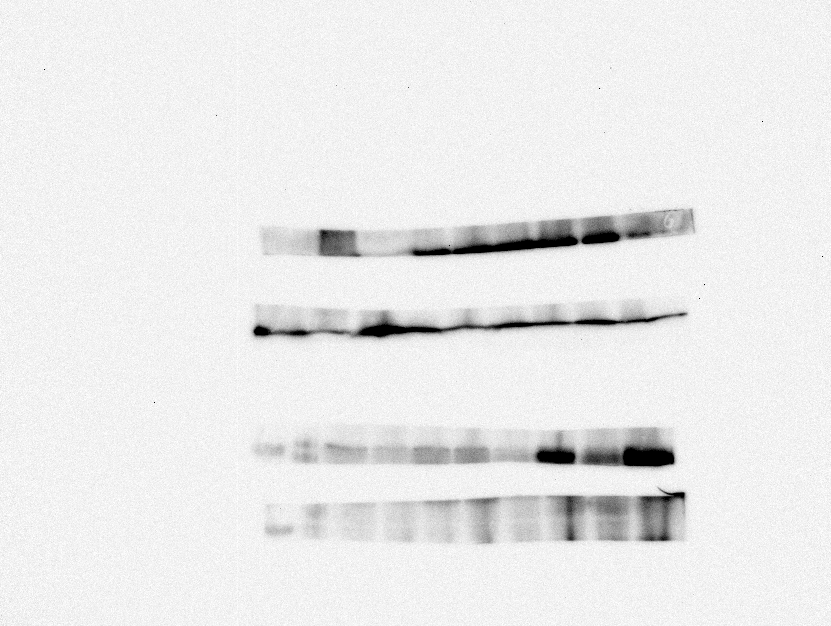

GAPDH

Figure 4: Unprocessed western blot for MYOCD, ANP and GAPDH proteins in MYOCD siRNA treated RAL rats (before and after ligation). Sequence from Left to right is: 1. Control rat 2. Renal artery ligated rat 3. Renal artery ligation + MYOCD silencing Pre-ligation 4. Renal artery ligation + MYOCD silencing Post-ligation 5. Renal artery ligation + Scrambled (negative siRNA)

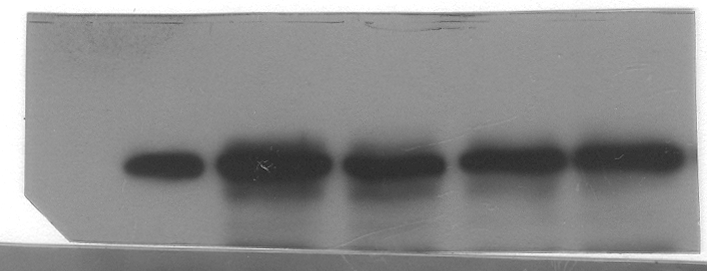

ANP

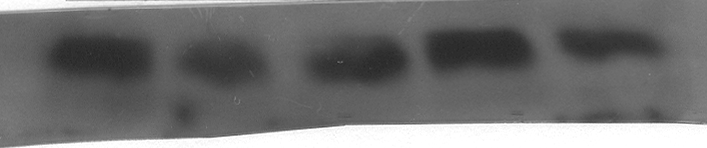

MYOCD

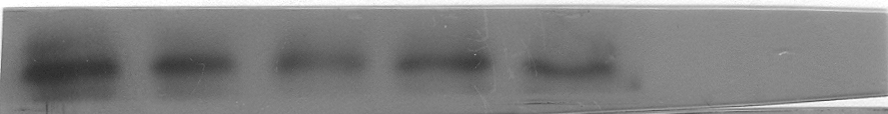

βMHC

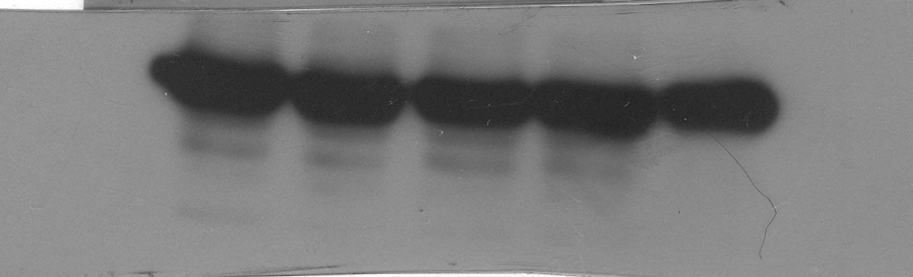

GAPDH

Figure 5: Unprocessed western blot for FGF, CTGF, MYOCD and GAPDH proteins in MYOCD siRNA treated RAL rats (before and after ligation). Sequence from Left to right is: 1. Control rat 2. Renal artery ligated 3. Renal artery ligation + Scrambled (negative siRNA) 4. Renal artery ligation + MYOCD silencing Pre-ligation 5. Renal artery ligation + MYOCD silencing Post-ligation

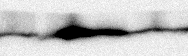

FGF

CTGF

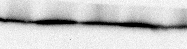

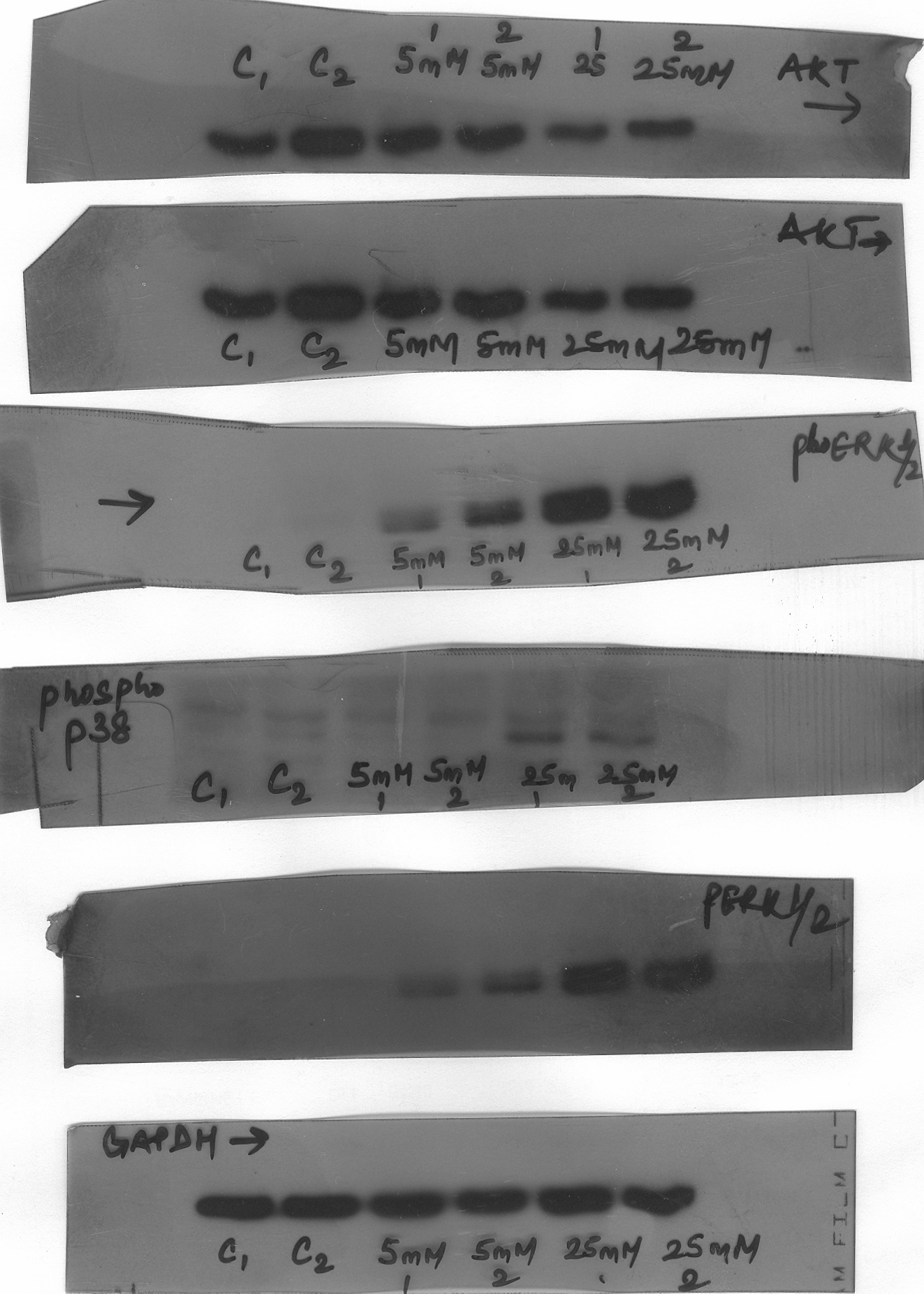

MYOCD

GAPDH

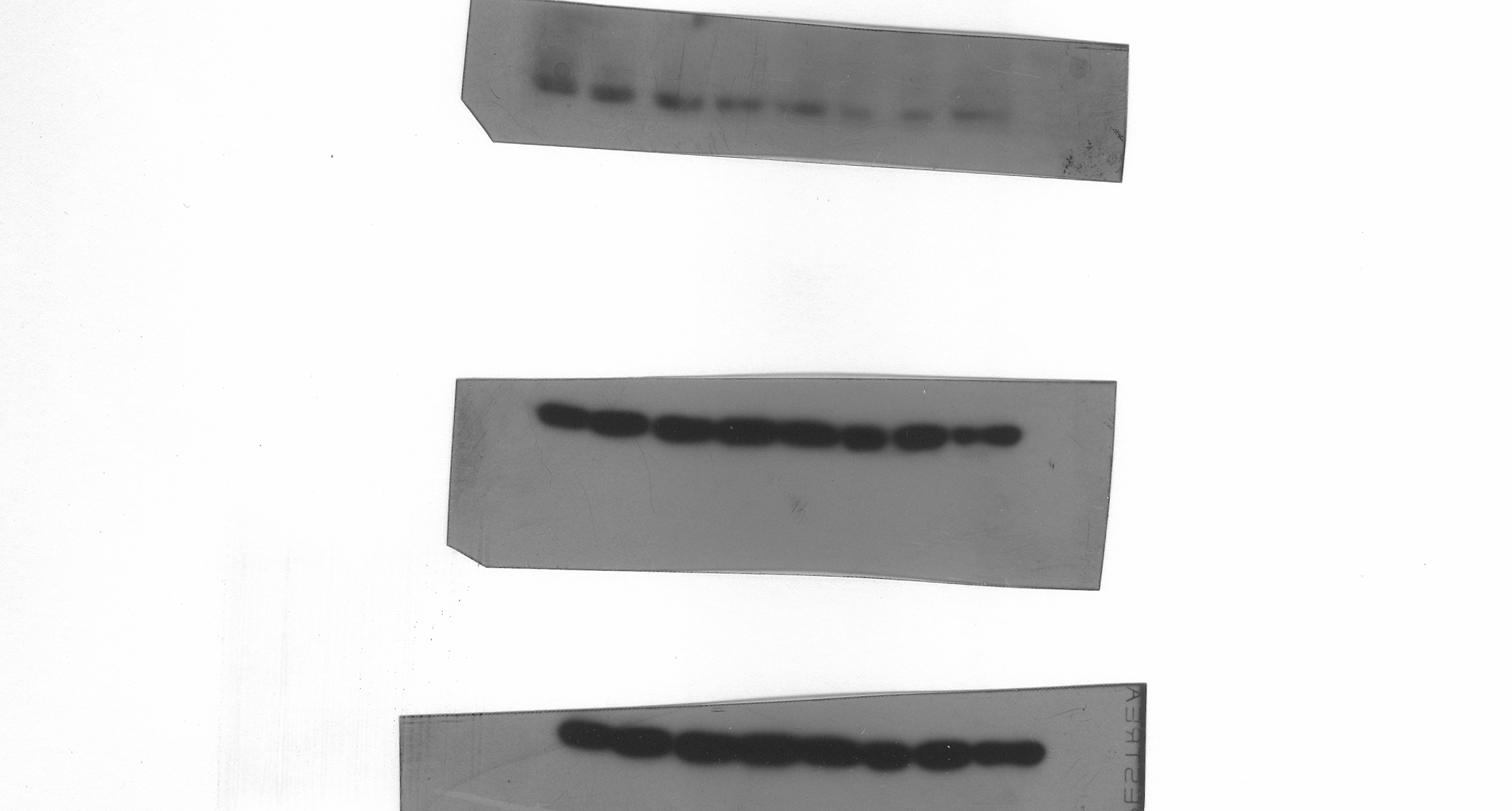

Figure 6a-d: H&E staining at lower magnification a) Control rat b) Renal artery ligated c) Renal artery ligation + MYOCD silencing Pre-ligation d) Renal artery ligation + MYOCD silencing Post-ligation. 6 e-h: MT staining at lower magnification e) Control rat f) Renal artery ligated g) Renal artery ligation + MYOCD silencing Pre-ligation h) Renal artery ligation + MYOCD silencing Post-ligation.

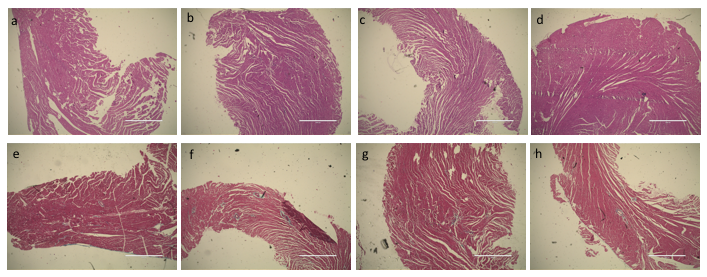
